# CATERPillar: A Flexible Framework for Generating White Matter Numerical Substrates with incorporated Glial Cells

**DOI:** 10.1101/2025.06.20.660694

**Authors:** Jasmine Nguyen-Duc, Malte Brammerloh, Melina Cherchali, Inès de Riedmatten, Jean-Baptiste Pérot, Jonathan Rafael-Patiño, Ileana O Jelescu

## Abstract

Monte Carlo diffusion simulations in numerical substrates are valuable for exploring the sensitivity and specificity of the diffusion MRI (dMRI) signal to realistic cell microstructure features. A crucial component of such simulations is the use of numerical phantoms that accurately represent the target tissue, which is in this case, cerebral white matter (WM). This study introduces CATERPillar (Computational Axonal Threading Engine for Realistic Proliferation), a novel method that simulates the mechanic of axonal growth using overlapping spheres as elementary units. CATERPillar facilitates parallel axon development while preventing collisions, offering user control over key structural parameters such as cellular density, undulation, beading and myelination. Its uniqueness lies in its ability to generate not only realistic axonal structures but also realistic glial cells, enhancing the biological fidelity of simulations. We showed that our grown substrates feature distributions of key morphological parameters that agree with those from histological studies. The structural realism of the astrocytic components was quantitatively validated using Sholl analysis. Furthermore, the time-dependent diffusion in the extra- and intra-axonal compartments accurately reflected expected characteristics of short-range disorder, as predicted by theoretical models. CATERPillar is open source and can be used to (a) develop new acquisition schemes that sensitise the MRI signal to unique tissue microstructure features, (b) test the accuracy of a broad range of analytical models, and (c) build a set of substrates to train machine learning models on.

**Highlights:**

- CATERPillar generates realistic synthetic voxels of white matter containing packed axons and glial cells.
- Synthetic axons had similar morphologies to those segmented from human electron microscopy in previous works.
- The morphological features of synthetic astrocytes closely matched those observed in histology.
- The functional form of diffusion and kurtosis time-dependence in the intra- and extra-axonal spaces agreed with experimentally observed disorder power laws in voxels composed of synthetic axons.

## 1. Introduction

Brain white matter (WM) consists of a dense network of axons originating from neurons whose cell bodies are mostly located in gray matter (GM). Many of these axons are encased in concentric layers of proteolipid membranes known as myelin, which serve to insulate them and enhance the speed and efficiency of electrochemical signal conduction (Flanagan et al., 2018). Axons serve as the communication gateway of the brain, allowing WM to function as the “edges” within the brain’s structural network (Zhang et al., 2021). Other cell types, such as glial cells, are also present in WM. These glial cells do not directly contribute to neuronal signalling, but perform essential support functions: oligodendrocytes promote myelination, astrocytes maintain the optimal chemical environment for neuronal communication and contribute to neurovascular coupling (Attwell et al., 2010), and microglia clear cellular debris from injury sites (Purves et al., 2001; Winther et al., 2024). Both axons and glial cells have highly dynamic morphologies and are influenced by their local environment (Andersson et al., 2020). To investigate the microstructure of these cells in WM, classical histology has proven to be fundamental to modern neuroscience as it reveals mechanisms of neural connectivity and communication, and exposes the cellular origins of neurodegenerative disease (Alexander et al., 2019). However, these methods are expensive, invasive, and destructive to the tissue, restricting their use to small samples from post-mortem specimens (Alexander et al., 2019). Nonetheless, thanks to their high level of anatomical detail, classical histological methods remain a reliable gold standard for tissue microstructure.

Diffusion magnetic resonance imaging (dMRI) has garnered interest within the scientific community as it shows potential for investigating brain tissue microstructure non-invasively. As the diffusion signals are sensitive to the random displacement of water molecules, we are able to study tissue on scales lower than image resolution (tens of micrometers vs millimeters) (Kiselev, 2017). The ability of dMRI to non-invasively capture features of intact tissue, combined with a broad field of view and relatively fast, costeffective acquisition, makes it particularly well-suited for in vivo brain studies (Alexander et al., 2019). The growing interest for accurate estimations of microstructural features from dMRI signals has driven the development of a wide range of analytical models (Alexander et al., 2019; Jelescu et al., 2020; Novikov et al., 2019; Stanisz et al., 1997). These models rely on simplifying assumptions to describe relevant geometric and diffusion-related characteristics. Nevertheless, the accuracy of these models and the robustness of their estimates to microstructural complexity and to deviations from the assumptions cannot be easily verified due to the absence of realistic ground truth (GT) microstructural parameters. New and robust validation approaches are thus essential.

One such approach is numerical simulations in synthetic substrates, which offer a flexible and cost-effective way to study the dMRI signal in realistic yet controlled environments. They elucidate a link between GT microstructural features and dMRI signals, in complex environments for which analytical solutions cannot be obtained (Fieremans and Lee, 2018). Briefly, to model water molecule trajectories and produce synthetic dMRI signals, a substrate representing the tissue of interest is first generated, after which Monte Carlo (MC) diffusion simulation tools (Yeh et al., 2013; Hall and Alexander, 2009; Rafael-Patino et al., 2020; Lee et al., 2021; Cottaar, 2022) are employed. This framework opens several possibilities: (i) to test and optimise the sensitivity of various diffusion acquisition schemes to tissue microstructure features for which no analytical formulation exists (e.g. branching, beading, and spines), (ii) to test the accuracy and robustness of a broad range of analytical models in substrate conditions that deviate from their assumptions, and (iii) to build a set of substrates to train machine learning models on, alleviating the need for analytical models altogether. Notably, numerical simulations have been previously utilised to explore DWI changes in response to dynamic physiological processes such as ischemic stroke (Budde and Frank, 2010) and neuronal activation (Spencer et al., 2024).

Various approaches for generating WM numerical substrates exist. These approaches either derive meshes directly from segmented histological data (Chin et al., 2002; Nguyen et al., 2018; Xu et al., 2018; Lee et al., 2020a; Abdollahzadeh et al., 2021) or comprise generative models that produce substrates “from scratch”, aiming to match GT tissue parameters derived from histology (Balls and Frank, 2009; Yeh et al., 2013; Ginsburger et al., 2019; Callaghan et al., 2020; Villarreal-Haro et al., 2023; Winther et al., 2024).

In the first case, the substrates are undeniably realistic, while the available configurations are constrained by specific histological samples. Amongst the frameworks that replicate WM tissue “from scratch” are MEDUSA (Ginsburger et al., 2019), CONFIG (Callaghan et al., 2020), CACTUS (Villarreal-Haro et al., 2023), and WMG (Winther et al., 2024). Each of these methods prioritises different aspects, such as computational efficiency, packing density and realism of the growth process. Furthermore, while some of these frameworks support the generation of glial cells (some more elaborate than others), their biological realism has never been evaluated. Such validation efforts have been exclusively focused on synthetic axons.

Here, we introduce a WM substrate generator entitled CATERPillar (Computational Axonal Threading Engine for Realistic Proliferation), which combines various strengths of previous generators. We go beyond the stateof-the-art by placing a strong emphasis not only on generating anatomically realistic axons and their myelin sheath, but also on incorporating biologically accurate glial cells. CATERPillar integrates the use of overlapping unit spheres from MEDUSA (Ginsburger et al., 2019), which are computationally efficient, and the natural fibre growth approach from CONFIG (Callaghan et al., 2020). As parallelised axonal growth is enabled, the run-time for substrate generation is considerably decreased compared to most previous generators. Additionally, CATERPillar provides precise control over axonal undulation and beading amplitude, while allowing to vary the number of fibre populations. A user-friendly GUI has been publicly released to make this novel tool accessible to all users, regardless of their programming experience. The code has also been released open-source so that users may modify it and/or suggest modifications which could improve this tool. The aim of this work is to introduce the methods behind CATERPillar and to validate the realism of the generated axons and glial cells (astrocytes in particular) by comparing quantitative features with EM-derived distributions, and the realism of the overall cellular layout. For the latter, the diffusion properties within each compartment were tested against theoretical predictions from effective medium theory (Novikov et al., 2014), confirming the short-range disorder behaviour expected for biological tissue, both inside the axons and in the extra-axonal space.

## 2. Methods

### 2.1. Growing WM substrates with CATERPillar

In the following, we first outline the main steps of the substrate growth algorithm and describe its implementation. Next, we provide a more detailed description of the creation of the different substrate constituents: glial cells, axons and myelin.

#### 2.1.1. Substrate growth implementation

The substrate is generated in a box-shaped volume through a multi-step growth process implemented in C++ programming language, following userdefined parameters governing the substrate growth (Figure 1). The substrate generation is performed as follows :

**Figure 1.**
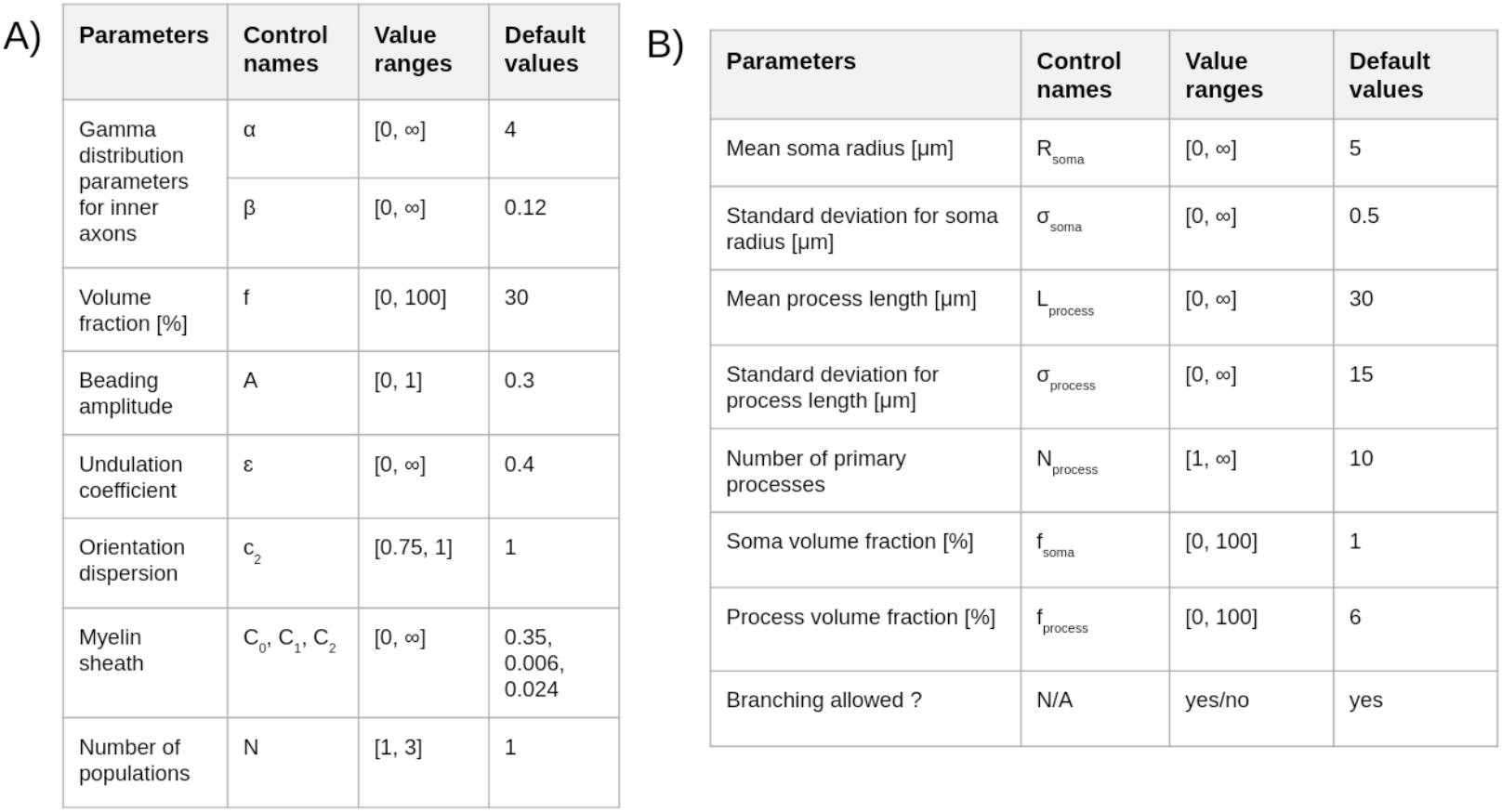
List of parameters used to generate populations of axons (A) and glial cells (B) in CATERPillar.

1. Glial cell somas are placed within the volume until a target volume fraction is achieved.
2. Axons are grown from one plane towards the opposite plane, with their trajectories shaped by parameters such as desired packing density, undulation, beading, and fibre orientation distribution function (fODF).
3. Glial processes extend from the somas, growing while avoiding collisions with existing axons, until the desired volume fraction is reached.

All cells are constructed by progressively placing overlapping spheres in chains. The spacing between two adjacent spheres is initially set to max(*R*_1_, *R*_2_) for efficiency, where *R*_1_ is the radius of the first sphere and *R*_2_ the radius for the next (see Supplementary Figure S1). To smooth the surface, additional overlapping spheres are added between consecutive ones, spaced by 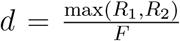, where *F* is a user-defined overlap factor. Higher *F* values improve realism but increase computational cost. Supplementary materials (Supplementary Figure S2) detail how *F* = 4 balances accuracy and computation time.

##### Development of the GUI

To improve the ease of use of the CATERPillar substrate generator, a GUI was developed in C++ using the Qt framework. It provides users with an intuitive and interactive environment to adjust the parameters that govern substrate generation. Additionally, the GUI supports the visualisation of the generated substrates, enabling users to inspect their spatial structure and composition. Figure 2 illustrates an example of a generated substrate containing both axons and astrocytes, showcasing the tool’s capability to render complex multicellular environments.

**Figure 2.**
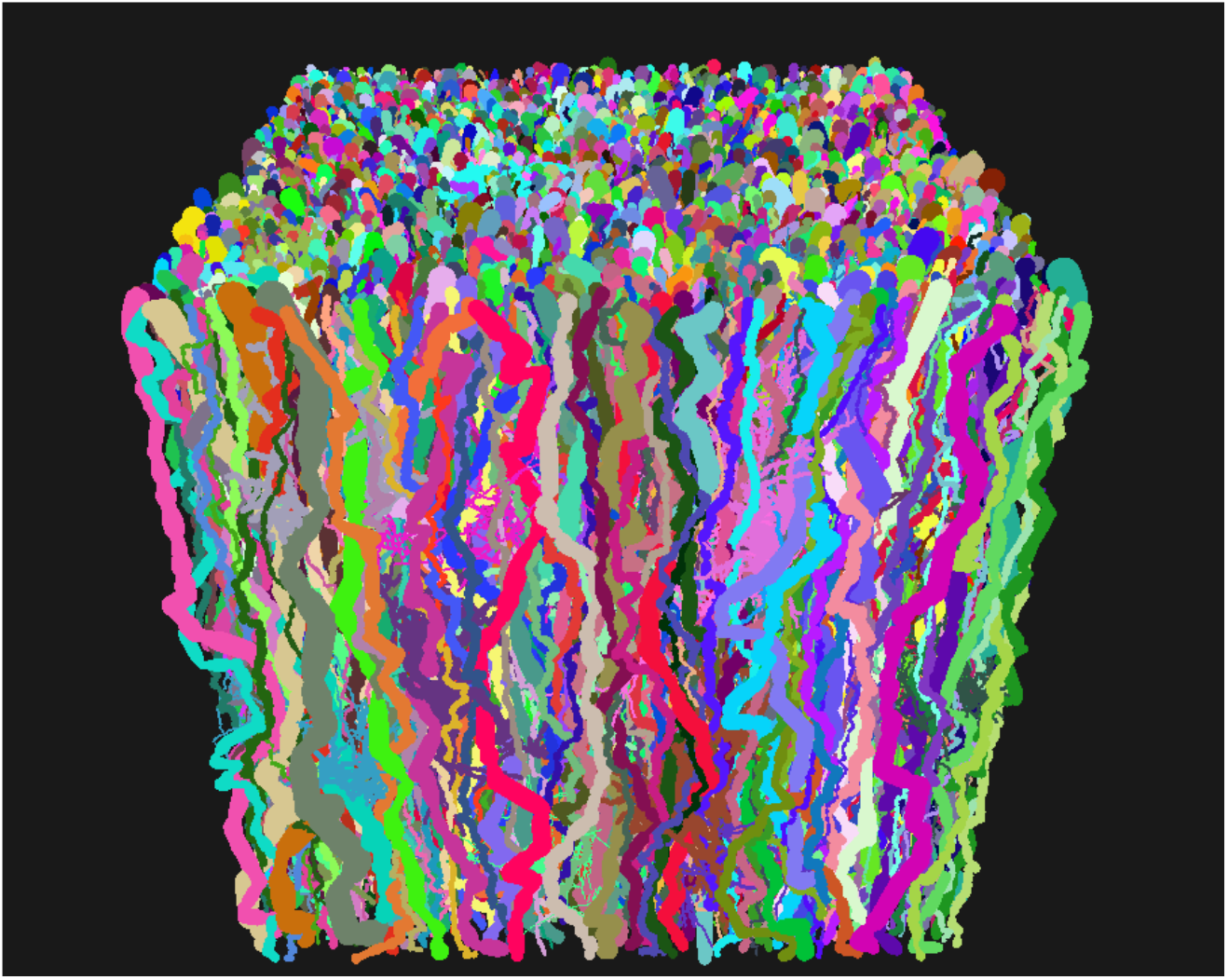
Example of a numerical substrate shown with the GUI’s visualisation tool.

#### 2.1.2. Axons

##### Initial growth process

The numerical substrate is a cube of edge length L. In order to generate axons in this cube, a set of radii are drawn from a representative Gamma distribution (user-defined by parameters *α* and *β*) (Assaf et al., 2008; Shekhar et al., 1996), until the ratio of their total disk areas to the voxel *x*-*y* plane area (*L*^2^) achieves the target density *f*. Each drawn radius is attributed to an axon, with a starting position placed randomly on a plane and an “attractor” on the opposite one. At each step of the growth process, successive spheres are sequentially positioned in a chain following the attractor’s direction. The axon grows until it reaches the opposite plane or collides with another obstacle. If an obstacle is encountered, multiple attempts are made to identify an alternative available position for the next sphere. If a suitable position cannot be found after several attempts, the radius of the sphere is gradually reduced to enable the axon to pass through tighter regions. This reduction typically affects only a small sequence of consecutive spheres and is therefore not expected to significantly alter the axon’s average radius. However, if growth remains obstructed despite these adjustments, the axon is deleted, and axon growth is reinitiated at a different starting position. The collision detection algorithm used is inspired by the Sweep and Prune method (Avril et al., 2011). This algorithm enhances efficiency compared to the brute-force approach by creating bounding boxes around each object. To boost the speed of the algorithm even further, we use threads to grow axons in parallel.

##### Control over undulation

During growth, the next sphere’s position is determined using polar and azimuthal angles, both sampled from Gaussian distributions centred at 0 to bias growth along the z-axis, before rotating toward the attractor. The standard deviation *ϵ*, which is user-defined, controls deviation from the attractor’s direction, and thus affects the axon’s undulation coefficient. The latter is defined as the total axon length divided by the straight-line distance between endpoints. Prior studies reported undulation coefficients around 1–1.2 for various axons (Stepanyants et al., 2004; Abdollahzadeh et al., 2021). We recommend *ϵ* = 0.4 to reproduce an undulation coefficient of 1.2. Figure 3A shows how *ϵ* influences undulations.

**Figure 3.**
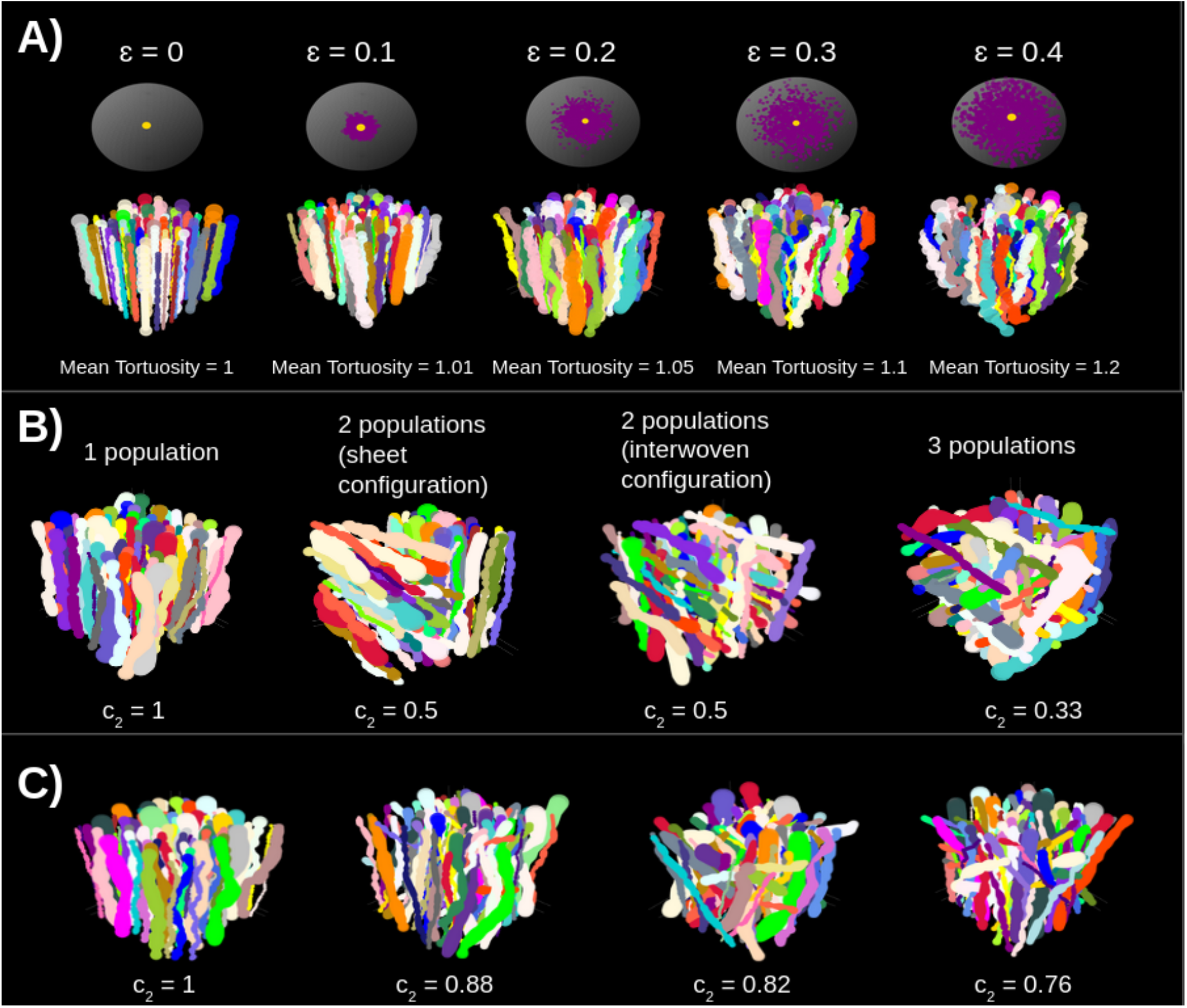
Illustrations of numerical substrates (width = 30 *µm*) with variations in specific parameters. A) Increasing the *ϵ* parameter leads to an increase in undulation values. B) Different number of fibre populations that cross each other perpendicularly. In case of 2 populations, they cross each other in two different ways: sheet configuration or interwoven configuration. C) The angles between the starting points of axons and their attractors can be set so that a specific orientation dispersion is obtained, leading to the desired *c*_2_.

##### Multiple fibre populations

Multiple fibre populations can be grown, with primary bundle directions perpendicular to one another. For two populations (N = 2), the user can specify whether the fibres grow as adjacent, parallel sheets or intersect perpendicularly in an interwoven pattern. For three populations (N = 3), the fibres are exclusively interwoven (see Figure 3B).

##### fODF

Users define the fibre orientation distribution using the parameter *c*_2_, representing the mean squared cosine of axon orientations relative to the primary bundle direction, assuming a Watson distribution. This parameter ranges from 1/3 to 1, with higher values yielding higher alignment of fibres. The *c*_2_ value is first converted into the corresponding Watson concentration parameter *κ*. Using the cumulative distribution function (CDF) table from (Zhang et al., 2011), which relates angles to *κ*, an orientation angle is sampled. This angle defines the deviation of the axonal growth direction from the main fibre orientation within the voxel. The CDF table provides data for *κ* values ranging from 4 to 128, which corresponds to a minimum achievable *c*_2_ of approximately 0.7. Achieving lower *c*_2_ values thus requires introducing additional fibre populations. Figure 3C shows examples of substrates with different *c*_2_ values.

##### Radius modulation

Axon radius variation is introduced stochastically to simulate random fluctuations along the axon length, while ensuring that the beading amplitude remains within a controlled range. Each new sphere’s radius (*R*_2_) is sampled from a Gaussian distribution centred on the previous radius (*R*_1_). Given the initially assigned radius for the axon (*R*_*i*_), if *R*_1_ falls outside the allowed range of *R*_*i*_ ± *A* · *R*_*i*_, the Gaussian distribution is recentred at the nearest boundary of this range. The parameter *A*, set by the user, is denoted the beading amplitude and controls the relative extent of radius fluctuations, and thus governs the degree of axonal beading.

#### 2.1.3. Myelin sheath

Once axonal growth is complete, myelin can be added by introducing spheres within the existing ones. This is achieved by duplicating each sphere and reducing its radius. The region between the inner and outer spheres constitutes the myelin compartment, whose thickness is determined by a user-specified g-ratio. This g-ratio 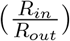 is calculated as follows, assuming that myelin thickness follows a log-linear relationship as previously established by Berthold et al. (1983) (Berthold et al., 1983):

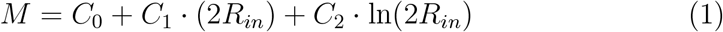

where M is the myelin thickness and *R*_*in*_ is the inner radius. A previous study using EM data (Lee et al., 2019) proposed a fitted model with parameters *C*_0_ = 0.35, *C*_1_ = 0.006, and *C*_2_ = 0.024 *µm*. In CATERPillar, users may specify the parameters *C*_0_, *C*_1_, and *C*_2_ (see Figure 1); however, we recommend keeping the default values to ensure that the generated myelin sheaths remain consistent with biologically observed structures. A lower bound of the g-ratio is imposed, ensuring that it does not fall below 0.2 for any sphere, in line with (Lee et al., 2019). The user may define and grow a population of myelinated axons and a population of unmyelinated axons, as the two may coexist within the same voxel.

#### 2.1.4. Glial cells

First, glial somas are placed in the volume using radii sampled from a user-defined Gaussian distribution (mean = *R*_*soma*_ and standard deviation = *σ*_*soma*_). Once the target for the glial soma volume fraction (*f*_*soma*_) is reached, axons grow while avoiding these somas, as previously described. Each soma then extends a user-defined number of primary processes (*N*_*process*_), defined as the branches originating directly from the soma. If branching is enabled, higher-order processes may sprout from existing ones until a target volume for processes (*f*_*process*_) is reached; otherwise, additional primary processes grow (see Figure 4). Processes grow in random directions, with their radii decaying exponentially with length. A decrease in the diameter of processes with their length was observed in microscopic images of glial cells in (Oberheim et al., 2009). We model this decrease as an exponential decay, as this was shown to be a good fit (Savtchenko et al., 2018). Growth continues until either a specific process length is reached or when an obstacle cannot be avoided. The length for each process is drawn from a Gaussian distribution with mean *L*_process_ and standard deviation *σ*_process_. It is defined here as the cumulative distance from the soma to the terminal tip, encompassing not only primary processes but also all higher-order branches. By measuring the full path from the soma rather than just individual branch segments, we ensure that higher-order processes are represented with proportionally shorter lengths compared to the primary processes.

**Figure 4.**
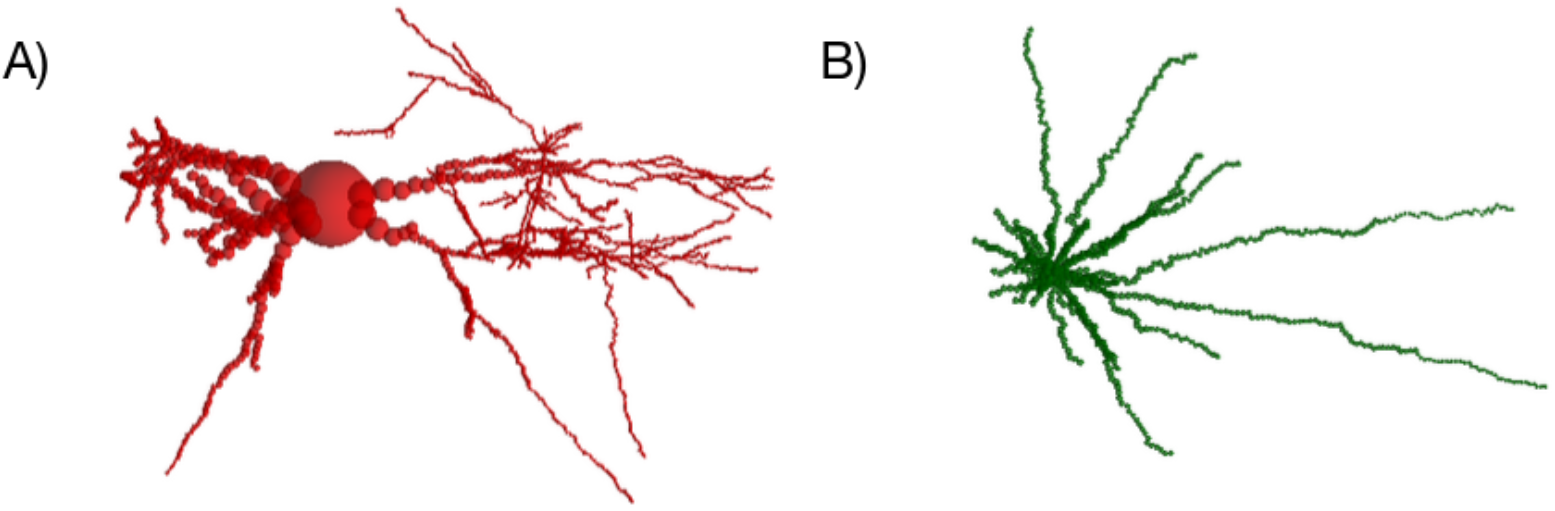
Example of (A) a synthetic glial cell with primary and secondary processes and (B) a synthetic glial cell with only primary processes. This example uses sparsely overlapping spheres to aid visual interpretation.

Our tool allows for generating up to two distinct glial cell populations within a single voxel. Each population can be independently configured with its own set of parameters. This enables the inclusion of various glial populations (such as astrocytes and oligodendrocytes) or the representation of healthy and pathological glial cells within the same substrate.

### 2.2. Morphological analysis

We analysed the morphology of both axons and glial cells in our numerical substrates. To evaluate the realism of myelinated axons, we generated a representative WM substrate with 1176 axons grown in a (100 *µ*m)^3^ voxel using default parameters shown in Figure 1A. Inner radii followed a Gamma distribution (*α* = 4, *β* = 0.12), and g-ratios were locally derived using a fitted model from (Lee et al., 2019). The beading amplitude was set to 0.3 and *ϵ* to 0.4. Resulting distributions for inner/outer diameters, g-ratio and diameter coefficients of variation (CV) were compared to previous findings on axons segmented from electron-microscopy, specifically to Figure 4 from (Lee et al., 2019).

For glial cells, substrates of (100 *µm*)^3^ were generated based on morphological parameters representative of rodent astrocytes, as detailed data for these cells are more widely available than their human counterpart (Reeves et al., 2011; Xin et al., 2019; Verkhratsky et al., 2018; Viana et al., 2023; He et al., 2024; Althammer et al., 2020; Klein et al., 2020; Köhler et al., 2023; Barsanti et al., 2024; Aird-Rossiter et al., 2025). These glial cells are thus specifically modelling astrocytes, so we will refer to them as such from now on. The parameters were set to the default values shown in Figure 1B. The analysis was performed on synthetic astrocytes grown in two settings: one within axon-filled substrates (79 astrocytes) with a volume fraction *f* of 30%, and another in empty space (64 astrocytes). As our synthetic astrocytes are expected to be adaptive to their environment (in this case, to the presence of axons), we expect to find morphological differences between the two groups, despite them having grown with the same set of parameter values. For clarity, we refer to these as protoplasmic and fibrous astrocytes, as these are the names given to astrocytes present within GM and WM tissue, respectively.

To demonstrate these differences, we applied Sholl analysis to quantify the arbour complexity of both astrocyte types by counting the number of intersections between astrocytic processes and concentric spheres centred on the soma. This Sholl analysis gave insight on the critical radius, process maximum and maximum radius, as illustrated in Figure 5. For each astrocyte, we also determined the total number of processes, the mean process radius, as well as the mean and total process length. This was done when taking into account only the primary processes, and for all processes combined. The statistical analysis between protoplasmic and fibrous astrocytes was conducted using the unpaired two-sample t-test in Python, and significance was assumed when *p <* 0.05. Additionally, we compared the structural fractional anisotropy (FA) that was computed as described in (Aird-Rossiter et al., 2025). Each process contributed to the FA of its corresponding astrocyte, with its influence weighted according to its volume. We expect fibrous astrocytes to exhibit greater elongation compared to those in empty space, resulting in higher FA values. By confirming this in our analysis, we also confirm that our cells are adaptive to their environment through their collision avoiding rules during growth.

**Figure 5.**
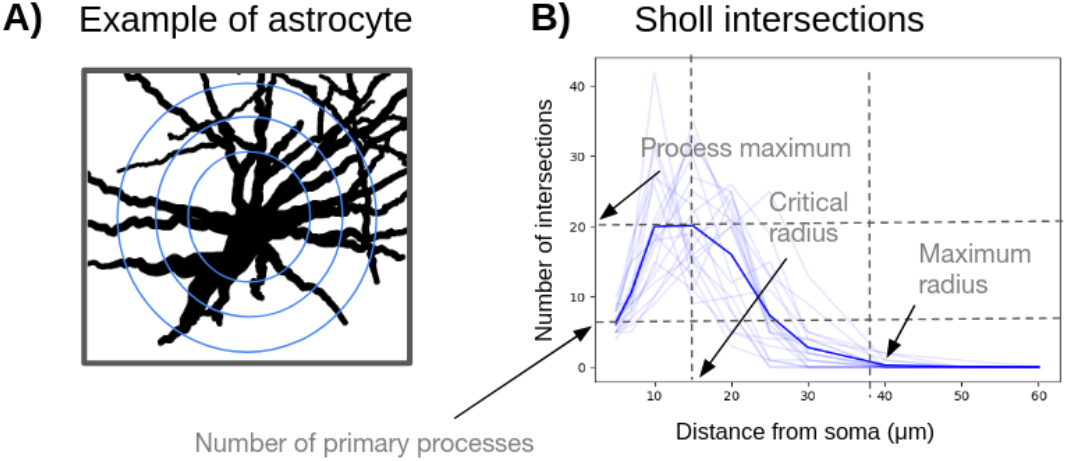
A) Illustration showing a synthetic astrocyte with concentric circles around its soma, depicting the concept behind Sholl analysis. B) In Sholl analysis, the process maximum refers to the highest number of intersections between astrocytic processes and the concentric circles. The critical value denotes the radial distance from the soma at which this maximum occurs, while the maximum radius corresponds to the furthest extent of the astrocytic processes (Baldwin et al., 2024). Individual Sholl intersection curves for each glial cell in a voxel (light lines) are averaged to produce the darker mean curve.

### 2.3. Monte Carlo simulations of water diffusion

After generating WM substrates and confirming the morphological realism of the different cell populations grown, we evaluated whether the signature of the dMRI signals simulated from synthetics voxels of packed axons reflected the expected patterns. To do so, we conducted MC simulations within various substrates containing impermeable unmyelinated axons and examined whether the time dependence of diffusivity in the intraand extraaxonal spaces aligned with theoretical predictions for short-range disorder from effective medium theory (Novikov et al., 2014). Before doing so, we adapted a previously published MC simulator to the specifics of the CATERPillar substrate generation.

#### 2.3.1. Adaptations and Extensions of the MC/DC tool

The diffusion simulator used in this work was the MC/DC (Monte Carlo Diffusion and Collision) simulator, a previously published open-source tool (Rafael-Patino et al., 2020). MC/DC integrates core dMRI protocols with a range of tissue models and supports various substrate geometries, including cylindrical, spherical, and mesh-based structures. Thanks to its opensource nature, MC/DC has been successfully adapted in multiple prior studies (Gardier et al., 2023; Villarreal-Haro et al., 2023). For the CEXI model (Gardier et al., 2023), the tool was extended to incorporate membrane permeability in permeable spherical cells.

For this study, we extended MC/DC to implement mirror boundary conditions. MC/DC originally relied on periodic tiling, where structures repeat seamlessly across boundaries. While effective for many scenarios, this approach is less suitable for substrates with high orientation dispersion. We introduced mirror boundary conditions, where random walkers cross voxel boundaries and diffuse within the mirrored substrate on the other side. Simulating diffusion within a mirrored environment has the advantage of maintaining microstructural continuity (Fieremans and Lee, 2018). However, it can increase the apparent fODF and introduce curvature artifacts. To minimise such edge effects, we placed walkers within a central region buffered from the edges by a distance of 30 *µm*. This distance should be at least equal to the mean squared displacement: ⟨ *x*^2^⟩ = 2*Dt*, where *D* is diffusivity and *t* is diffusion time. Additionally, to support intracellular diffusion, we modified MC/DC to allow walkers to diffuse across overlapping spheres by ignoring collisions at surfaces that lie within neighbouring spheres.

#### 2.3.2. Time dependence of diffusivity

Numerous studies (Burcaw et al., 2015; Fieremans et al., 2016; Jespersen et al., 2018; Lee et al., 2020a,b, 2021, 2025) have explored the structural disorder (Novikov et al., 2014) within biological tissues and have shown that the time dependence of diffusion and kurtosis can be modelled according to the structural universality class assigned to the medium. It has been shown that brain tissue is best characterised by the short-range disorder class.

To verify that the CATERPillar-generated substrates display characteristics of this short-range disorder, Diffusion Kurtosis Imaging (DKI) was computed using the TMI toolbox (Novikov et al., 2018; Coelho et al., 2022; Ades-Aron, 2018) across multiple diffusion times, separately for the intraaxonal space (IAS) and extra-axonal space (EAS). Theoretically, if the substrate exhibits short-range disorder as expected, the axial diffusivity (AD) and kurtosis (AK) should scale as 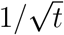, while the radial diffusivity (RD) and kurtosis (RK) are expected to follow a ln(*t/δ*)*/t* dependence (Burcaw et al., 2015; Fieremans et al., 2016; Jespersen et al., 2018; Lee et al., 2020b,a, 2021). The value *δ* refers to the PGSE pulse width and *t* is the diffusion time. More specifically, we expect :

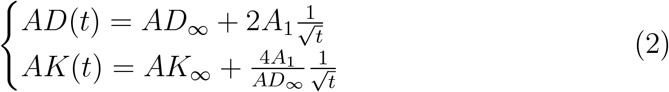

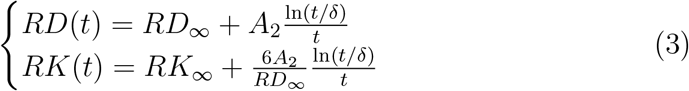

In the radial direction, these relationships remain valid as long as *t* ≫ *δ*, while in the axial direction, they hold provided that *t* ≫ *t*_*c*_, where *t*_*c*_ represents the characteristic diffusion time across the disorder correlation length *l*_*c*_. This *l*_*c*_ was typically reported within the range 1-20 *µm* (Fieremans et al., 2016) *depending on the structure of the WM tissue and on the compartment and direction of interest. Determining l*_*c*_ and *t*_*c*_ is non trivial, but assuming the 1-20 *µm* range, we can assume that *t* ≫ *t*_*c*_ and *t* ≫ *δ* is valid when *t >* 20*ms*.

The reduction in diffusivity described over time, by the model, is in theory driven by the increasing number of interactions between water molecules and cellular membranes (Novikov et al., 2014). These collisions limit the mean squared displacement of the molecules, thereby lowering the diffusion. In contrast, kurtosis increases at short diffusion times before decreasing at longer times, with the latter trend being captured by the model (Lee et al., 2025). For further details on disorder classes, refer to the Supplementary Materials 13.2.

To assess the time dependence of the diffusion and kurtosis, numerical simulations were performed on substrates of 150 *µ*m per side. The axonal volume fraction was approximately 50%, and the axons were unmyelinated yet impermeable, in a single bundle, without glial cells. Three types of substrates were generated: substrate with axons modelled as cylinders (*S*1), one with beaded axons with a beading amplitude of 0.3 (*S*2), one with undulating (*ϵ* = 0.4) and beaded axons (*S*3). The radii of axons were drawn from a Gamma distribution with *α* = 4 and *β* = 0.25.

Simulations used bulk diffusivities of 2.5 *µ*m^2^/ms for IAS and 1.5 *µ*m^2^/ms for EAS. Random walkers were initialised in a central (90 *µ*m)^3^ region within the (150 *µ*m)^3^ cube to avoid edge effects. A step size of 0.1 *µ*m yielded 115,500 and 69,300 steps for IAS and EAS, respectively. Simulations used 10^5^ walkers per compartment (proven to be sufficient in (Hall and Alexander, 2009)), repeated three times per substrate to capture variability.

The analysis focused on diffusion times of 11, 16, 25, 44, 109 and 157 ms as similarly done in (Lee et al., 2020a). To approximate a narrow-pulse Pulse Gradient Spin Echo (PGSE) sequence, the gradient duration *δ* was fixed at 4 ms, while the inter-gradient duration Δ and gradient strength *G* varied to obtain different combinations of diffusion times and b-values. Sixty unique gradient directions per shell were generated with optimal angular coverage using the method outlined by (Caruyer et al., 2013). The b-values utilised were 0, 500, 1000 and 2000 *s/mm*^2^. Diffusion and kurtosis tensors were independently estimated from the simulated IAS and EAS signals. From these, AD, RD, AK, and RK were derived for each compartment and analysed as functions of diffusion time. The metrics were fitted to the model described in equations 2 and 3, using only diffusion times above 20 ms to ensure that *t* ≫ *t*_*c*_. The resulting fit was then plotted across the full range of diffusion times.

## 3. Results

### 3.1. Performance of WM substrate generation with CATERPillar

Figure 6 shows the time necessary to grow a substrate of 150 *µm* in length for various axonal packings, when setting *ϵ* to 0.4 and the beading amplitude to 0.3. By parallelising the axonal growth computations across 20 threads in this example, high packing densities (up to 70%) could be achieved for this voxel size under 12 hours. Each thread used one core and approximately 300 MB of RAM.

**Figure 6.**
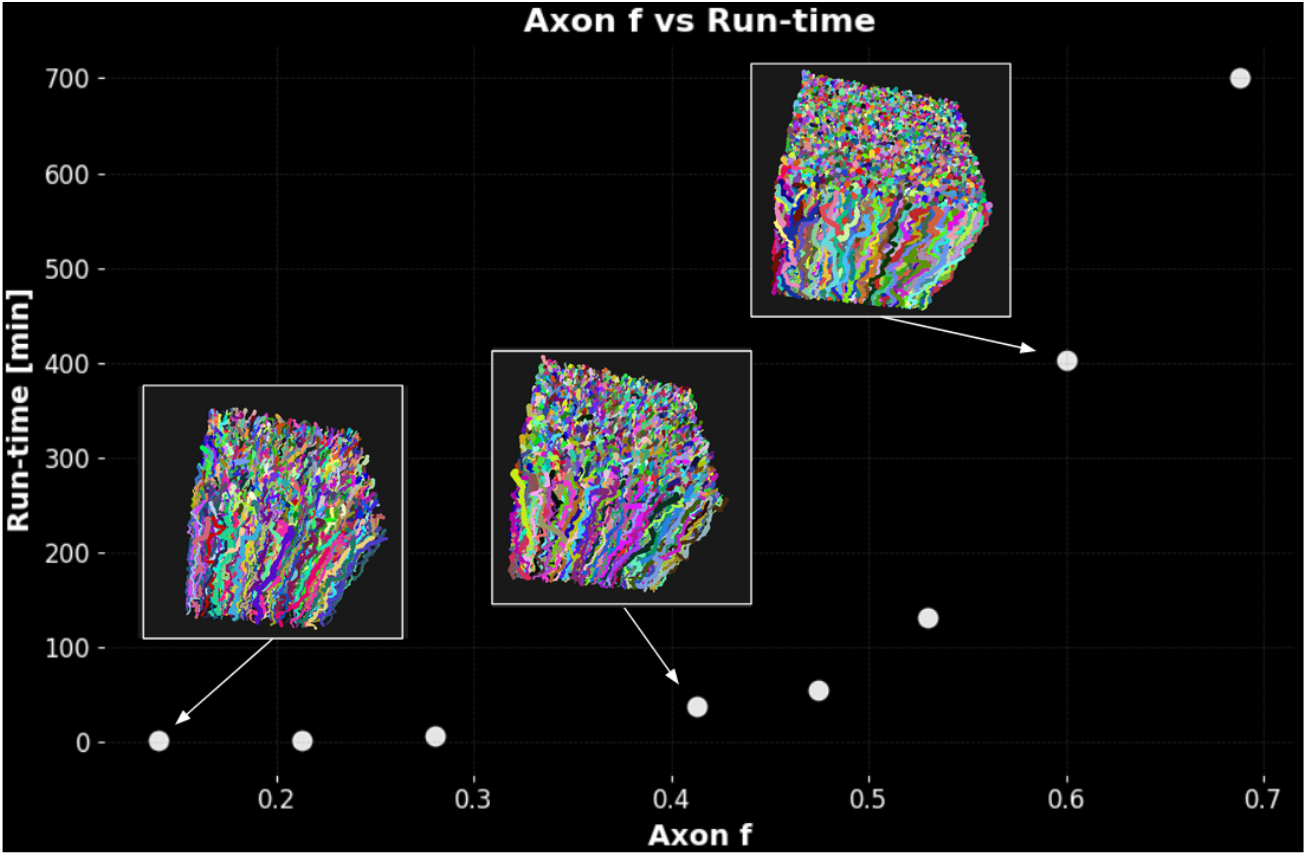
The run-time versus the axonal packing *f* for substrates of 150 *µm* in length.

### 3.2. Morphological analysis

#### 3.2.1. Myelinated Axons

We evaluated our synthetic axon morphologies against histological measurements illustrated in (Lee et al., 2019) (see Fig. 4 in that work), and the results were highly consistent. Figure 7 shows histograms of inner and outer diameters, g-ratios, diameter CV, and undulation for our synthetic axons. The near-perfect alignment of these distributions with those in Figure 4 from (Lee et al., 2019) indicated that our generation pipeline faithfully reproduced real axonal geometry. Specifically, our g-ratios tracked the expected inner-diameter dependence from Equation 1, staying between 0.2 and 1, and the CV peaked around 0.2 for outer diameters and 0.3 for inner diameters.

**Figure 7.**
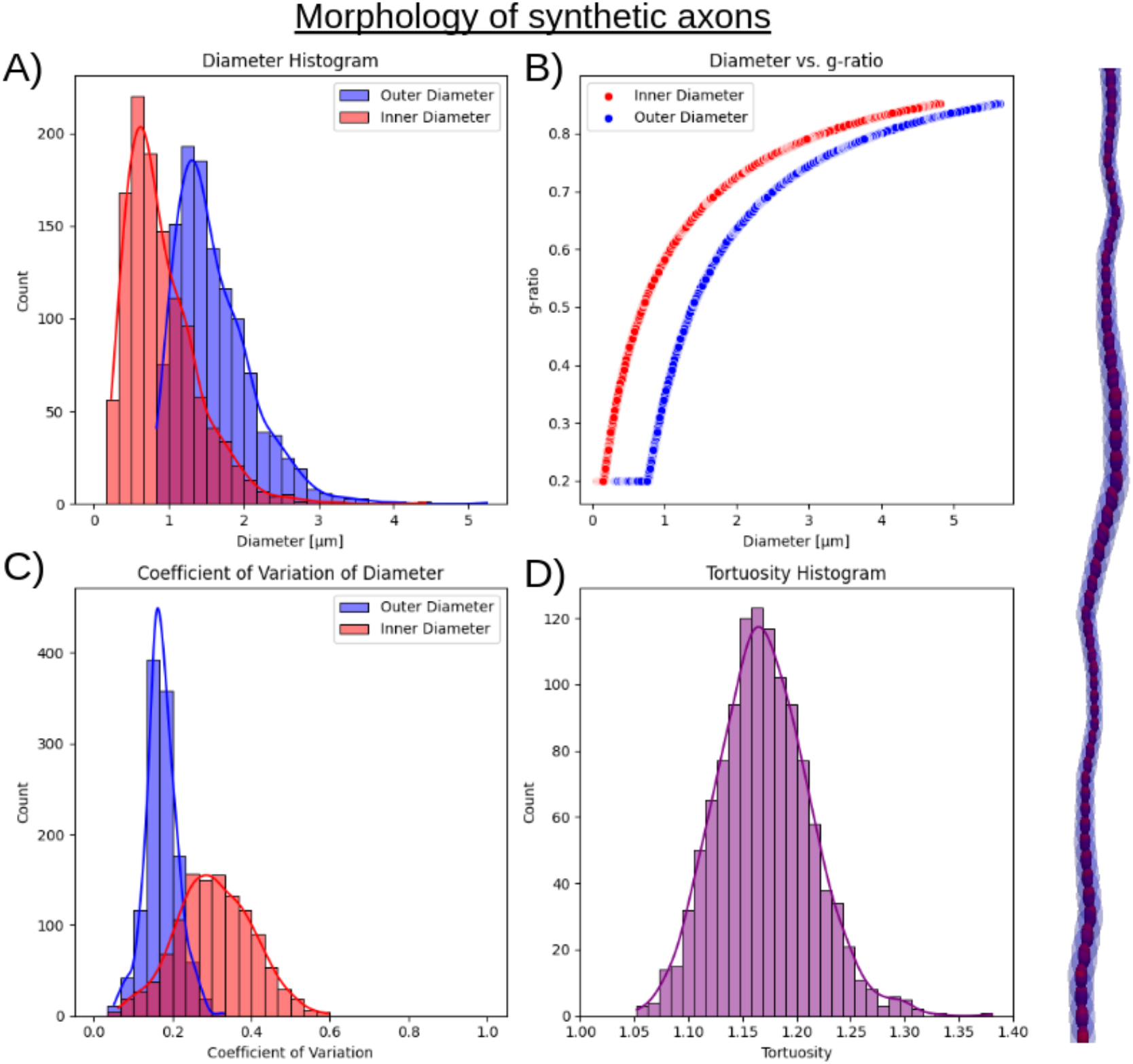
Several histograms showcasing the morphology of synthetic axons: A) Histogram of the inner (red) and outer (blue) diameters of approximately 1176 myelinated axons within a voxel of 100 *µm* in length. The Gamma distribution parameters (for inner radius distribution) were *α* = 4 and *β* = 0.12. B) Histogram of the g-ratios of these axons with respect to the inner and outer diameter. This follows the log-linear relationship fit from (Lee et al., 2019). C) Histogram of the Coefficients of Variation for inner and outer diameters, resulting from a beading amplitude of 0.3. D) Histogram of tortuosities, obtained using *ϵ* = 0.4.

Undulation, which was not reported in (Lee et al., 2019), followed an almost Gaussian profile centred at 1.18 (range 1.05–1.4), matching physiological values reported by (Stepanyants et al., 2004). These close correspondences confirmed that our CATERPillar-generated substrates effectively captured the realistic structure of WM axons.

#### 3.2.2. Astrocytes

Figure 8A and B show the two types of astrocyte-containing substrates: one with astrocytes grown in empty space and the other with astrocytes grown in a voxel populated with axons. The averaged Sholl intersection curves in panel C, computed across all astrocytes within each voxel, reflected their characteristic arbour complexity. The curve for both synthetic protoplasmic and fibrous astrocytes closely matched previous histological findings in rodent GM (Reeves et al., 2011; Tavares et al., 2017; Viana et al., 2023; Althammer et al., 2020; Klein et al., 2020; Barsanti et al., 2024; He et al., 2024), showing the typical increase in intersections up to a process maximum between 20 and 50 intersections (y-axis), at a critical radius between 20 and 30 *µm* (x-axis), followed by a gradual decline ending between 40 and 60 *µm*. Fibrous astrocytes exhibited a higher maximum process length than protoplasmic astrocytes, while their critical radius was slightly smaller.

**Figure 8.**
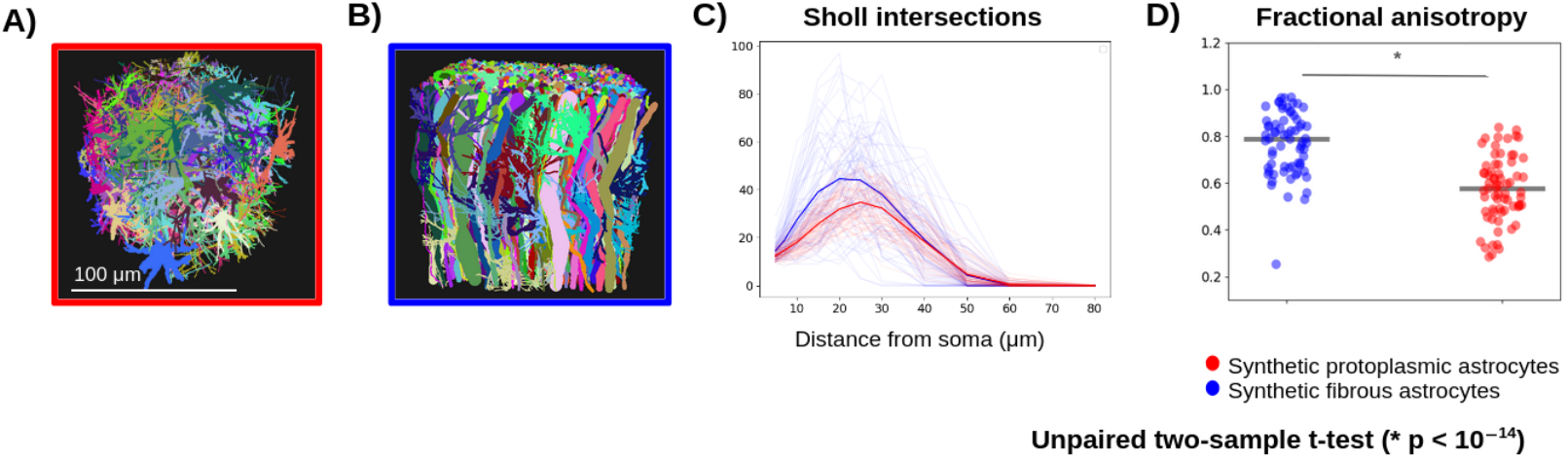
A) Illustration of synthetic astrocytes grown in empty space. The voxel is 100 *µm* in length. B) Illustration of synthetic astrocytes grown in a voxel already filled with axons (*f* = 30%). C) Sholl analysis of the two kinds of astrocytes, in red for those shown in A and in blue for those in B. D) Differences in FA values between the two groups, with each scattered point referring to a specific astrocyte. Statistical significance was obtained using the unpaired two-sample t-test.

In Figure 9, further morphological analysis examined the number of processes, total process length, mean process length, and mean process radius for both astrocyte types. These measurements were assessed separately for primary processes (panels A-D) and for all processes (panels E-H). These results agreed well with previous histological findings obtained from protoplasmic astrocytes in rodents, illustrated with a pink boxes (Tavares et al., 2017; Viana et al., 2023; Barsanti et al., 2024; Klein et al., 2020; He et al., 2024; Aird-Rossiter et al., 2025). In panel A, both astrocyte types exhibited a median close to 10 primary processes, consistent with the input specification. In panel C, the median primary process length for both astrocyte types fell below the 30 *µm* input target, reflecting possible premature terminations of growth when collisions with obstacles occurred. Although we did not explicitly control the features shown in panels B, C, F, and G, they nonetheless remained within the expected ranges derived from rodent histological protoplasmic data. Panel D demonstrates that the mean radius of primary processes exceeded that of all processes (panel H), consistent with the fact that process radius decreased as length increases. Overall, the morphological differences indicated that fibrous astrocytes possessed a greater number of processes that were, on average, shorter than those of protoplasmic astrocytes.

**Figure 9.**
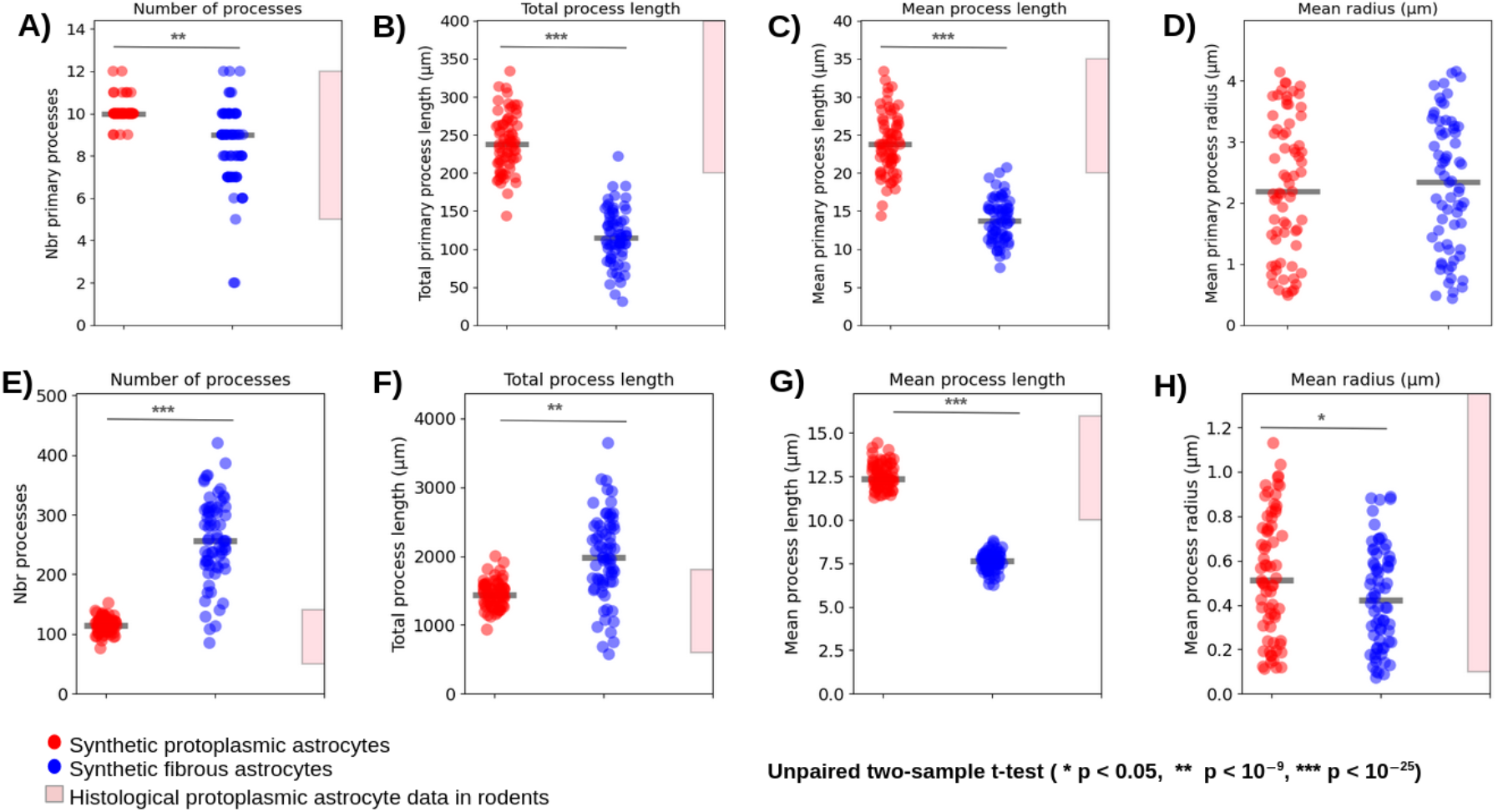
Number of processes, total process length, mean process length and mean process radius shown when taking into account A) only the primary processes, B) all processes. This is shown for synthetic protoplasmic and fibrous astrocytes. Each scattered point refers to an astrocyte. The pink boxes show the typical ranges for each morphological feature in histological findings for protoplasmic astrocytes in rodent GM (Tavares et al., 2017; Viana et al., 2023; Barsanti et al., 2024; Klein et al., 2020; He et al., 2024; AirdRossiter et al., 2025). Statistical significance was obtained using the unpaired two-sample t-test.

### 3.3. Time dependence of diffusivity

To assess whether the structural disorder of the generated substrates fell within the short-range class, the time dependence of the diffusion D(t) and the kurtosis K(t) was assessed. This was done within the IAS and the EAS separately, in the axial and radial directions and across the three different substrate configurations, each composed of impermeable unmyelinated axons (Figure 10). For simplicity, we will refer to the substrate containing cylinders as *S*1, the substrate containing beaded axons as *S*2 and the last one containing undulating and beaded axons as *S*3.

**Figure 10.**
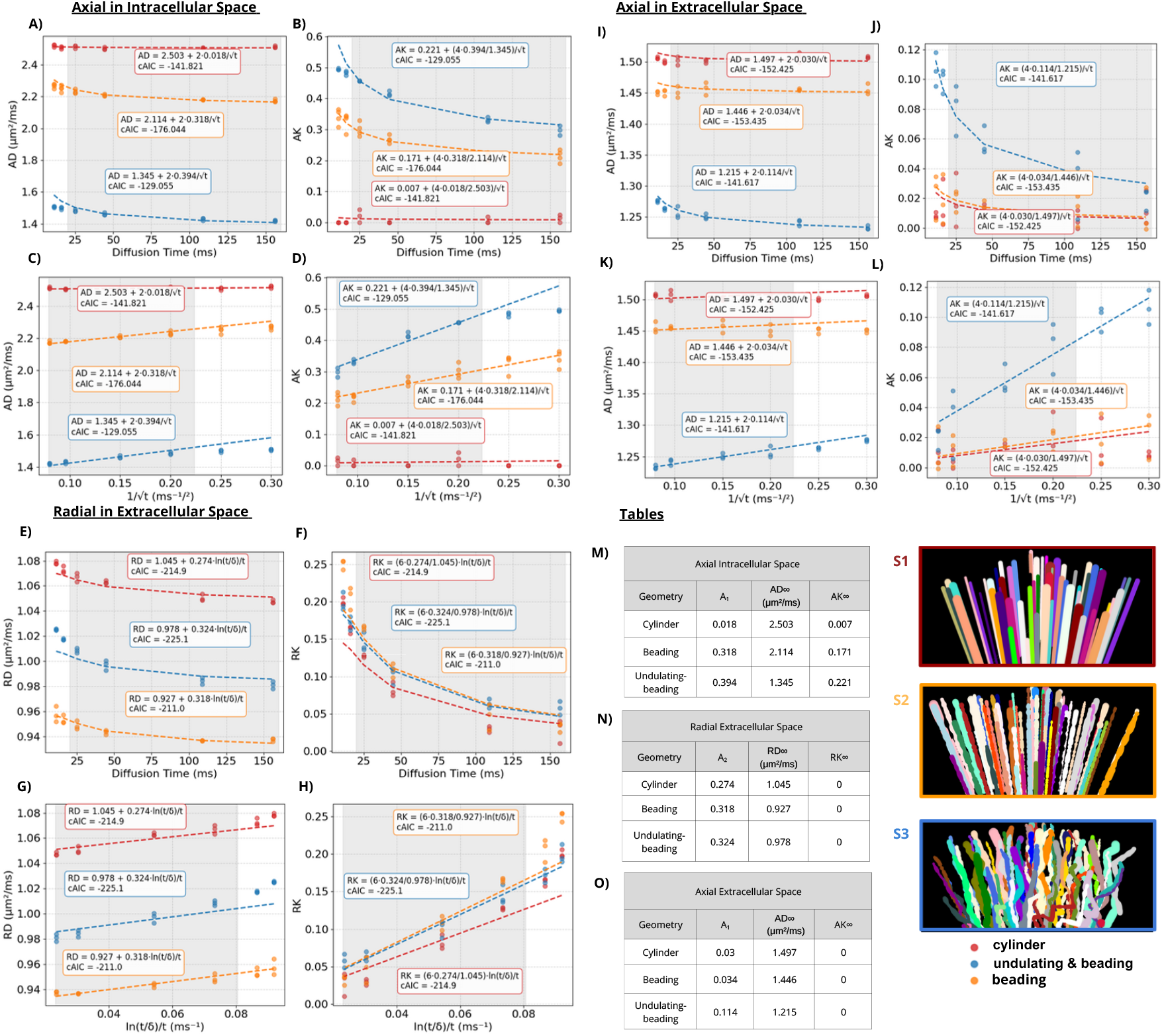
Time dependence of diffusion and kurtosis. A–D) Axial diffusivity and kurtosis in the IAS were fit jointly, with corresponding fitted parameters shown in M. E–H) Radial diffusivity and kurtosis in the EAS were fit jointly with fitted parameters in N. I–L) Axial diffusivity and kurtosis in the EAS were fit jointly with fitted parameters in O. All fits were performed using equations 2 and 3. The grey region marks the range of diffusion times used for model fitting. Colours represent different substrates (*S*1, *S*2 and *S*3). Each simulation was repeated three times to assess variability.

Within the IAS (Figure 10A-D and M), both AD and AK remained constant for *S*1. AD was approximately equal to the free diffusivity assigned to the IAS (2.5 *µ*m^2^/ms), and AK was equal to 0. In contrast, *S*2 and *S*3 showed time dependence of AD and AK, with a good agreement with the 1D short-range disorder model. The long-time asymptotic diffusivity *AD*_∞_ was higher for *S*2 than for *S*3 (2.11 and 1.35 *µ*m^2^/ms), whereas the value for *A*_1_ was lower for *S*2 than for *S*3 (0.32 and 0.39 respectively). The *AK*_∞_ values were non-zero for both *S*2 and *S*3.

As shown in Figure 10E–H and N, the extracellular RD and RK exhibited time dependence across all substrates, in good agreement with the 2D shortrange disorder model that predicted a *ln*(*t/δ*)*/t* dependence. The long-time asymptotic diffusivity *RD*_∞_ was 1.05, 0.93, and 0.98 *µ*m^2^/ms for substrates *S*1, *S*2, and *S*3, respectively, reflecting the effective extra-axonal tortuosity in each substrate. The parameter *A*_2_ was similar across all substrates but slightly increased with axonal structural complexity (ranging from 0.27 to 0.32).

The AD within the EAS, as shown in Figure 10I–L and O, remained constant (close to the free diffusivity value assigned to this compartment 1.5 *µ*m^2^/ms) across diffusion times for both *S*1 and *S*2, while AK was nearly zero, with some weak time-dependence at short times. In contrast, *S*3 displayed lower AD than the free diffusivity value, positive AK, and marked time-dependence of both AD and AK, which followed functional form characteristic of 1D disorder.

Collectively, the corrected Akaike Information Criterion (cAIC) values were below -100 for all substrates and conditions, indicating a very good fit to the respective models for all conditions. Indeed, the trends at long diffusion times agree well with the expected functional forms, but sometimes deviate at short times, where the long-time limit does not apply.

## 4. Discussion

We present CATERPillar, a new tool for generating numerical WM substrates that is inspired by the use of overlapping spheres (as in MEDUSA (Ginsburger et al., 2019)) and of natural fibre growth (from CONFIG (Callaghan et al., 2020)). The use of overlapping spheres improves computational efficiency, as collision detection between walkers and cell membranes is faster with spheres than with triangular mesh representations (Ginsburger et al., 2019). Contextual growth, on the other hand, more closely mimics natural fibre development compared to applying repulsion forces to pre-existing structures, which may be important for generating more realistic phantoms (Callaghan et al., 2020). Beyond the combination of these two assets, and unlike earlier tools focused mainly on axons, CATERPillar realistically models both axons and glial cells based on histological data. It supports the creation of beaded, undulating, and dispersed axons (with or without myelin) and multiple glial populations, making it suitable for simulating physiologically relevant conditions and validating microstructural models. Voxel sizes ranged from 100–150 *µm* in this study, though bigger voxels may be obtained at the cost of a longer run-time. The current packing limit is approximately 70%, and the maximum dispersion of fibres is *c*_2_ = 0.75 when only one fibre population is employed. Three crossing populations can achieve a *c*_2_ value for dispersion as low as 1/3.

### 4.1. Run-time

Creating numerical substrates of 150 *µm* in length and with 50 % packing took approximately 2 hours when using 20 cores, each using approximately 300 MB of RAM. For substrates of 70% packing, this run-time reached 12 hours. A comparison with the CACTUS tool remained difficult, as the runtime published was for a 90% packing in very large voxels (500 *µm*). In this case, the optimisation ran on 24 cores (each using 400 MB of RAM) and completed in 4 hours, while the reconstruction was performed on a single core, taking approximately 1 minute per fibre for a total of 33,478 fibres (Villarreal-Haro et al., 2023).

Compared to CONFIG and WMG, CATERPillar showed improved runtime performance: CONFIG required approximately 6 hours and 9.4 GB of RAM, while WMG took around 31 hours with unspecified memory usage to generate a cubic voxel of about 50 *µm* in size. In terms of fibre packing, WMG achieved 72% and CONFIG reached 75%, both comparable to CATERPillar’s maximum packing of 70% (Callaghan et al., 2020; Winther et al., 2024).

However, CATERPillar remained much slower than the GPU - based MEDUSA, which could generate 100 *µm* voxels with a packing of 65% in just 56 seconds. This is partly attributable to our use of a growth-based model instead of repulsion-based packing, which is more computationally efficient and allows for denser arrangements. Additionally, the use of CPU significantly slows substrate generation compared to GPU-based approaches. Future releases aim to improve runtime by implementing CATERPillar to be run on GPUs.

### 4.2. Realism of Synthetic Axons

#### 4.2.1. Morphological analysis

To validate axonal realism, we compared diameter distribution, g-ratio scaling, beading and undulation with distribution obtained from histological data (Lee et al., 2019). We found strong agreements that support the biological accuracy of the generated axons. To obtain such accuracy, we used *ϵ* = 0.4 for undulation, beading amplitude = 0.3 and the Gamma distribution parameters (for inner radius distribution) were *α* = 4 and *β* = 0.12. If the user wishes to grow unmyelinated axons, we would advise a Gamma distribution with parameters *α* = 4 and *β* = 0.25, as this should be close to the corresponding outer radius distribution. Further adjustments can be made by the user if a specific WM region is targeted for example, and the axons are known to have different properties there.

#### 4.2.2. Time dependence

Beyond morphology, we evaluated the time dependence of the diffusivity coefficient and kurtosis metrics in three substrates with unmyelinated impermeable axons. The substrates featured axons with varying levels of structural complexity, ranging from simple cylinders in *S*1, representing the least realistic model, to *S*3, which most closely resembled real WM by incorporating both undulations and beading. Because real WM tissue exhibits short-range structural disorder, theoretical models predict that AD and AK should follow a 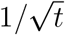 dependence, while RD and RK are expected to scale with *ln*(*t/δ*)*/t* (Novikov et al., 2014; Burcaw et al., 2015; Fieremans et al., 2016). Moreover, the time-dependent behaviours of diffusion and kurtosis are intrinsically linked, as they share common parameters in the models described by equations 2 and 3. Additionally to obtaining a good fit for this model, we expect the structural complexity of the substrates to affect the resulting fitted parameters such that a higher structural complexity would induce lower diffusivities and higher kurtosis, as previously reported (Şimşek et al., 2025).

Overall, the fit was strong across all three conditions (EAS/axial, IAS/axial, EAS/radial) and for all three substrates (*S*1, *S*2, *S*3), as reflected by very low cAIC values. Deviations between the fit and the measurements could be noticed when *t <* 20*ms* (not included in the fit), as the model does not hold within these short times. Some additional minor deviations could also be observed when *t >* 20*ms*, but were very small considering that the model was fitted jointly for diffusivity and kurtosis.

When examining the strength of time dependence captured by *A*_1_ and *A*_2_, a consistent trend emerged across all three conditions: the values increased with the structural complexity of the substrate, being lowest for *S*1 and highest for *S*3. This trend was expected, as more irregular microstructural features better approximate biological tissue.

In the axial direction, *A*_1_ revealed distinct roles for the intraand extraaxonal spaces. *A*_1_ within the IAS appeared primarily sensitive to axonal beading, while within the EAS, it rather captured the effects of axonal undulations. This interpretation is supported by the absence of time dependence in *S*1’s IAS (where axons were cylindrical) and by the very weak dependence observed in the EAS for *S*1 and *S*2, where axons lacked undulation. The sensitivity of AD in the IAS to beading is consistent with previous findings (Lee et al., 2020a). To our knowledge, this is the first demonstration that axial diffusivity and kurtosis in the EAS can capture signatures of axonal undulation. Beading had little effect on *A*_1_ in the EAS, as shown by the minimal differences found between *S*1 and *S*2. However, in more densely packed substrates, beading could exert a stronger influence by increasing water-membrane interactions.

As for the radial direction of the EAS, both the diffusion and kurtosis exhibited time dependence across all substrates, with non-zero *A*_2_ values. This was expected, as the axons are positioned randomly in all substrates.

Regarding the long-time asymptotic diffusivities (*AD*_∞_ or *RD*_∞_), these metrics theoretically capture the medium’s tortuosity, being the extent to which diffusion is hindered by complex, non-linear pathways compared to free, straight-line motion. This tortuosity value can be described by the relationship *ς* = *D*_0_*/D*_∞_ − 1 (Lee et al., 2020b). Thus, when *D*_0_ = *D*_∞_, tortuosity *ς* is zero and an unrestricted diffusion is observed. In our results in the axial direction, higher tortuosity (lower *AD*_∞_) consistently corresponded to stronger time dependence (higher *A*_1_). This higher tortuosity was observed in more structurally complex substrates, the highest being for *S*3 and the lowest for *S*1.

Surprisingly, in the EAS, *RD*_∞_ suggested greater tortuosity in *S*2 than in *S*3. This outcome may seem counter-intuitive, as *S*3 is structurally more complex due to the presence of undulations. A possible explanation is that *S*2 has slightly higher packing density or that beading alone creates local pockets that hinder diffusion in the EAS more than when axons are also undulating.

Interestingly, long-time asymptotic kurtosis values (*AK*_∞_ or *RK*_∞_) were only observed to be greater than zero in the axial direction of the IAS. This may reflect inter-axon heterogeneity in the IAS as water molecules in different axons have distinct environments, so the overall kurtosis does not vanish. In contrast, in the EAS, all water resides in a single shared medium, leading to *AK*_∞_ = *RK*_∞_ = 0 (Lee et al., 2020b).

In summary, we examined how diffusion and kurtosis evolve with diffusion time across substrates of varying structural complexity. The short-range disorder model was jointly fitted to diffusion and kurtosis data, and the resulting parameters provided insights into both the strength of time dependence and the medium’s tortuosity, which matched the theoretical predictions and contribute to establishing the reliability of the grown synthetic voxels. This conclusion is further supported by a complementary analysis of the two-point correlation function presented in the Supplementary Materials (Supplementary Figure S3). Among all substrates, *S*3 was the only one to consistently exhibit substantial time dependence characterised by short-range disorder, and tortuosity across all conditions, which makes it the substrate closest to biological tissue.

### 4.3. Realism of Synthetic Astrocytes

Astrocytic morphology was assessed in two conditions: substrates containing only astrocytes, and substrates where astrocytes grew in the presence of axons. We hypothesised that astrocytes grown in axon-rich environments will adopt a more elongated morphology compared to those generated in empty space, leading to higher FA values. This elongation would reflect the typical morphology of fibrous astrocytes found in real WM tissue, in contrast to the more isotropic shape of protoplasmic astrocytes commonly located in GM (Köhler et al., 2023; Palombo et al., 2017). Also, we expected morphological characteristics of our synthetic astrocytes to align with histological observations from rodent protoplasmic astrocytes, as their growth parameters were defined based on such data.

As anticipated, the FA of synthetic fibrous astrocytes was significantly higher than that of protoplasmic astrocytes. This confirms that, through simple rules conducting collision avoidance during process growth, we were successful in making our astrocytes adaptive to their morphology to their environment. Regarding the additional morphological characteristics, those obtained from synthetic protoplasmic astrocytes fell within the ranges reported in previous histological studies of rodent protoplasmic astrocytes, confirming that they exhibited realistic arbour complexity. This held for both user-defined features and those that were not explicitly controlled. The morphological characteristics of fibrous astrocytes differed from those of protoplasmic astrocytes, and were sometimes slightly outside of the expected range for histological protoplasmic astrocytes. These differences reflected the shorter and more numerous processes of fibrous astrocytes, likely due to growth being constrained by frequent collisions with surrounding axons. In real tissue, fibrous astrocytes are known to be have a larger size, less branched and straighter processes, and fewer fine processes than protoplasmic astrocytes, which contrasts with our results. However, it is important to note that the same set of parameters were applied for both substrates to specifically demonstrate the impact of the environment on astrocyte morphology. Also, these parameters were set to specific values to match histological findings on protoplasmic astrocytes in rodents. To properly model fibrous astrocytes, the *L*_*process*_ would need to be much higher than the one used in this study. Keeping *f*_*soma*_ and *f*_*process*_ constant, the increase of *L*_*process*_ should reduce the need for branching, and thus decrease the number of processes.

## 5. Limitations

The main limitation of the CATERPillar tool lies in its achievable axonal volume fraction, which reaches approximately 70%, notably lower than the ∼90% reported with the CACTUS tool (Villarreal-Haro et al., 2023). Unlike CACTUS, which uses an optimisation algorithm to resolve initial fibre overlaps, CATERPillar models axonal growth by extending fibres from an initial point toward an attractor, aiming to emulate natural development as previously done in CONFIG. While this biologically inspired approach may enhance realism, it also increases computational demands and inherently limits the maximum packing density. Nonetheless, a packing fraction of 70% is likely a realistic estimate, aligning with histological observations that typically report around 80% intracellular and 20% extracellular space (Syková and Nicholson, 2008). It is worth noting, however, that chemical fixation procedures tend to shrink the extracellular space, potentially leading to an overestimation of intracellular volume fractions in histological data (Korogod et al., 2015).

Additionally, in terms of axonal morphology, it has been noted that axonal cross-sections are not always perfectly round but can be elliptical in shape. A study using EM-based segmentation reported an average eccentricity of 0.72 for myelinated axons in the contralateral corpus callosum (Abdollahzadeh et al., 2021), indicating a notably elongated cross-sectional shape, as a circle would have an eccentricity of 0. While this elliptical crosssection was accounted for in the CACTUS (Villarreal-Haro et al., 2023), CONFIG (Callaghan et al., 2020) and WMG tools (Winther et al., 2024), it had not been explicitly incorporated into CATERPillar. Nonetheless, the cross-section of our substrates in the x-y plane seemed to naturally adopt elliptical shapes due to the undulation of the axons, but the resulting eccentricity was not assessed. The effective signature of elliptical cross sections on the dMRI signal remains to be clarified.

## 6. Conclusions and Future Works

The CATERPillar framework facilitates the generation of WM numerical substrates, incorporating axons, myelin, and glial cells. It enables efficient fibre growth toward predefined attractors, allowing multiple axons to develop simultaneously. Additionally, CATERPillar provides precise control over key morphological parameters, such as undulation and axonal beading. This tool can be used to (a) develop new acquisition schemes that sensitise the MRI signal to unique tissue microstructure features, (b) test the accuracy of a broad range of analytical models, and (c) build a set of substrates to train machine learning models on. Through this study, we have demonstrated that the generated substrates accurately replicate axonal and astrocytic morphological characteristics observed in histological studies, reinforcing the tool’s capability for realistic WM modelling. The resulting morphology yielded time-dependent diffusivities consistent with experimental observations of 1D and 2D structural disorder. Hence, we will provide an open-source software for creating realistic synthetic WM substrates, which will help advancing the capabilities of dMRI to quantify WM microstructure in vivo by training numerical models of complex features instead of analytical models of simple geometrical features.

## 8. Code availability statement

The GitHub code and GUI for CATERPillar has been made public and can be found at: https://github.com/jazz031195/CATERPillar.

The Monte Carlo Simulator, adapted from (Rafael-Patino et al., 2020), can be found at: https://github.com/jazz031195/Permeable_MCDS/tree/whitematter.

## 9. Author Contribution

Conceptualization: I.J, J.N.D; Methodology : I.J, J.N.D, M.C, M.B; Software : J.N.D, M.C, J.R.P; Validation : J.N.D, M.B; Formal analysis: J.N.D; Writing-original draft: J.N.D; Writing - Review & Editing: I.J, J.N.D, M.B, J.P, I.d.R, J.R.P; Visualisation: J.N.D, M.B; Supervision: I.J.;

## 10. Acknowledgements

We are grateful to Rita Oliveira for testing the CATERPillar tool in her study, which contributed to improving its usability. This work was supported by the Swiss Secretariat for Research and Innovation (SERI) under an ERC Starting Grant award ‘FIREPATH’ MB22.00032. I.J. is supported by an SNSF Eccellenza Fellowship no. 194260.

## 11. Declaration of generative AI and AI-assisted technologies in the writing process

During the preparation of this work the author used ChatGPT 4o in order to rephrase certain sentences for improved readability. After using this tool/service, the author reviewed and edited the content as needed and takes full responsibility for the content of the publication.

