## Supplementary material for "CATERPillar: A Flexible Framework for Generating White Matter Numerical Substrates with incorporated Glial Cells": Supplemetary Materials

### Supplementary Materials

#### 1. Overlapping distance between unit spheres

To evaluate the effect of the spacing between overlapping spheres on diffusivity metrics, we compared synthetic DWI signals from straight axons modelled with varying sphere distances to those obtained using a perfect cylindrical representation. The objective was to identify the optimal $F$ value that minimises discrepancies from the cylindrical model while balancing computational efficiency, as runtime increases with higher $F$ values. To achieve this, we examined results for $F = 1, 2, 4$, and 8.

Substrates with dimensions $(150\,\mu m)^3$ containing straight axons were generated, one for each specified value of $F$, plus an additional substrate where axons were represented as cylinders. The IAS volume fraction was consistently maintained at 50%. Simulations featured $10^5$ random walker trajectories, with nine independent runs for each substrate type. A step size of $0.1\,\mu m$ was employed, while IAS and EAS diffusivities were set at $2\ \mu m^2/ms$. The selected PGSE sequence used a wide pulse, defined by $\Delta = 55.5\,\text{ms}$ and $\delta = 16.5\,\text{ms}$. DWIs were acquired along the x, y, and z axes, and apparent diffusion was calculated using the formula $-\frac{\log(DWI_2/DWI_1)}{b_2-b_1}$, with $b_1 = 0$ and $b_2 = 1000\,\text{ms}/\mu m^2$. RD was determined by measuring the apparent diffusion in the transverse direction to the axons (y-axis), while AD was calculated based on the apparent diffusion along the axonal orientation (z-axis). An unpaired two-sample t-test was applied to measure significant differences between the diffusivity metrics obtained with different values of $F$.

The degree of overlap between the spheres, as expected, influenced both radial and axial diffusivity in the IAS and the EAS. As shown in Figure 2B, decreasing the distance between the spheres (or increasing the value of $F$) caused the diffusivity values to more closely resemble those measured in a cylindrical substrate. In other words, higher $F$ values reduced the deviation from the idealised diffusivity associated with cylinders. However, this improvement in accuracy came with a trade-off: as illustrated in Figure 2C,

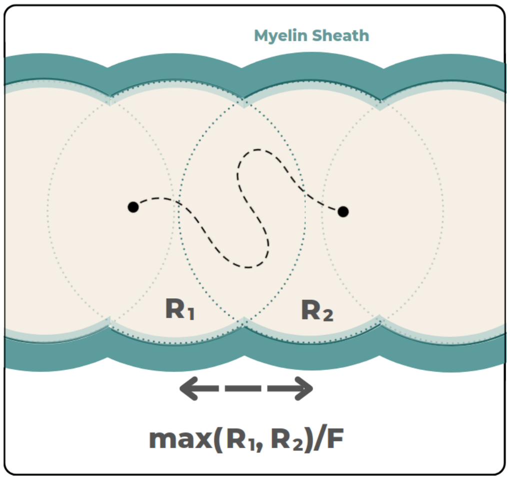


Figure 1: Illustration depicting how $F$ is related to the distance between consecutive spheres and their inner radius.

increasing $F$ led to a steep rise in computational run-time, following an exponential trend.

The selected $F$ value should therefore balance computational efficiency with reliable diffusivity results. Based on the data, $F = 4$ emerged as a suitable compromise. With this choice, the computed diffusivity was sufficiently close to that of cylinders, while keeping the run-time manageable. Consequently, a value of $F = 4$ was adopted for the analyses on the grown substrates.

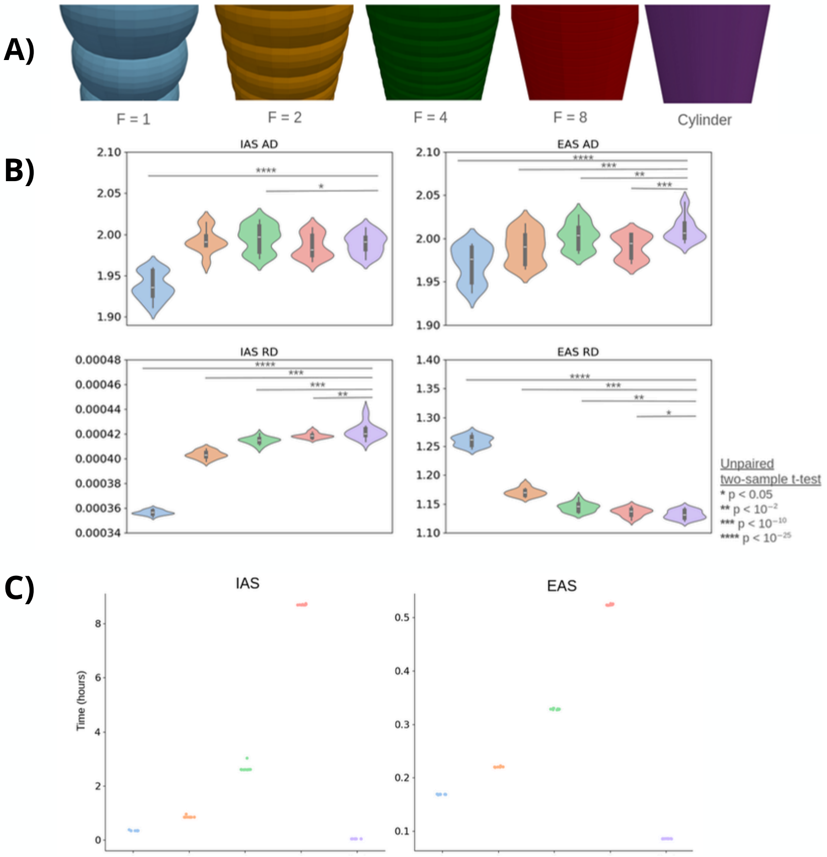


Figure 2: (A) Illustration of axonal segments demonstrating the influence of the factor $F$ on morphology. The spacing between two spheres is determined by the maximum radius divided by $F$. (B) DKI-derived diffusivity estimates for the IAS and EAS across different $F$ values. A trend of convergence is observed as $F$ increases, with the highest $F$ value yielding diffusivity estimates closest to those obtained from substrates where axons are modelled as cylinders. (C) Computational run-time for each type of substrate, showing an increasing trend with $F$, highlighting the trade-off between morphological accuracy and computational efficiency.

#### 2. Effective medium theory

The dMRI signal reflects the collective behaviour of a large population of diffusing spins. In conventional q-space imaging performed at a fixed diffusion time (Callaghan et al., 1991), much of the microstructural detail is obscured due to the averaging effect imposed by the macroscopic size of imaging voxels (Fieremans et al., 2016). In contrast, acquiring data across multiple diffusion times provides a means to probe structural restrictions at the scale of the diffusion length, which is within the micrometer scale (Lee et al., 2020b). Thus, the time dependence of the diffusion $D(t) =< x >^2 /2t$ and of the kurtosis $K(t) =< x^4 > / < x^2 >^2 -3$ (Kiselev, 2010; Jensen and Helpern, 2010) have been explored in previous studies (Latour et al., 1994; Barazany et al., 2009; Burcaw et al., 2015; Fieremans et al., 2016; De Santis et al., 2016; Lee et al., 2018, 2020a) to infer properties of the underlying microstructure. The structural disorder of the tissue has also been assessed, as it can also be quantified through the behaviour of $D(t)$ and $K(t)$ as a function of diffusion time.

According to effective medium theory (Novikov et al., 2014), the degree of this disorder is governed by the long-range spatial correlations of the restrictions (e.g. the cell membranes in biological tissue). Traditionally, this has been characterised using the two-point correlation function $\Gamma(\mathbf{k})$, derived from the Fourier transform of a binary mask representing the medium. When the wave vector k is sufficiently low ($k \to 0$), then $\Gamma(\mathbf{k})$ behaves like $k^p$ (Burcaw et al., 2015), with the $p$ constant being the structural exponent. Determining the value of $p$ allows the medium to be classified into a specific universality class, providing insight into the degree of structural disorder within the underlying tissue.

Several universality classes have been identified. One of these classes is called "Order" and corresponds to periodic arrangements with $p = \infty$. Another class, which is the most commonly observed, is called "Short-Range Disorder" and is associated with $p = 0$. The "Hyperuniform Disorder" class represents systems with $p > 0$, while the "Extended Disorder" class corresponds to $p < 0$. Previous works suggests that the most appropriate universality class for describing tissue structure is the short-range disorder (Burcaw et al., 2015; Fieremans et al., 2016; Jespersen et al., 2018; Lee et al., 2020b,a, 2021). Such findings were obtained with Monte Carlo simulations conducted on axons segmented from EM data (Lee et al., 2020a, 2021, 2020b), in experimental phantoms composed of parallel fibres designed to mimic axons (Burcaw et al., 2015) as well as in *in vivo* studies (Jespersen et al., 2018; Fieremans et al., 2016; Lee et al., 2020b). Since this class is associated with $p = 0$, we expect $\Gamma(\mathbf{k})$ to have a finite plateau in the low-$q$ regime ($q^0 = const$). This indicates the absence of long-range correlations beyond the correlation length $l_c \sim \frac{1}{k_c}$, with $k_c$ being the extent of this low-$k$ plateau (Burcaw et al., 2015). Physically, $l_c$ represents the characteristic distance from a given restriction beyond which positional information about other restrictions is lost, effectively erasing any "memory" of their locations (Fieremans et al., 2016).

As discussed earlier, the diffusion $D(t)$ and the kurtosis $K(t)$ each exhibit a specific time-dependent behaviour depending on the structural universality class corresponding to the medium. These dynamics are governed by the structural exponent $p$ and the spatial dimensionality $d$, through the derived exponent $\theta$, defined as:

$$\theta = \frac{p+d}{2}$$

The value of $\theta$ determines the temporal decay of diffusion and kurtosis according to the following general expressions (Lee et al., 2020b):

$$D(t) = \begin{cases} D_\infty + c_D\, t^{-\theta}, & \theta < 1 \\ D_\infty + A_2 \dfrac{\ln(t/\tilde{t}_c)}{t}, & \theta = 1 \\ D_\infty + \dfrac{\text{const}}{t}, & \theta > 1 \end{cases} \tag{1}$$

$$K(t) = \begin{cases} K_\infty + c_K\, t^{-\theta}, & \theta < 1 \\ K_\infty + \xi(p,d)\, \dfrac{A_2}{D_\infty}\, \dfrac{\ln(t/\tilde{t}_c)}{t}, & \theta = 1 \\ K_\infty + \dfrac{\text{const}}{t}, & \theta > 1 \end{cases} \tag{2}$$

In these equations:

- $D_\infty$ and $K_\infty$ are the long-time limits of diffusion and kurtosis.
- When $\theta < 1$, $c_D = \frac{A_1}{1-\theta}$
- $c_D$, $c_K$, $A_1$, and $A_2$ are constants characterising the structural features.

- $\tilde{t}_c = \max(t_c, \delta)$, where $t_c$ is the correlation time related to the structural correlation length $l_c$ and $\delta$ is the PGSE pulse width.

- $t >> t_c = \frac{l_c^2}{2D_\infty}$

The dimensionless function $\xi(p, d)$ relates the decay of kurtosis to that of diffusion and is given by (Lee et al., 2020b):

$$\xi(p,d) = \frac{c_K}{c_D/D_\infty} = 6\left[\left(2 + \frac{p(3p+d-4)}{2(d+2)}\right)\frac{1}{2-\theta} - 1\right]$$

Here, $d = 1$ corresponds to axial metrics (along neurites or axons), and $d = 2$ corresponds to radial metrics.

When $\theta < 1$, equations 1 and 2 rely on the narrow pulse approximation (with $\delta = 4$ ms in our case). If this condition is not met, corrections must be applied to account for the finite pulse width, as described in (Lee et al., 2021). For the case $\theta = 1$, determining the appropriate value of $\tilde{t}_c = \max(t_c, \delta)$ is more complex, as $t_c$ is generally unknown. To address this, we treated $t_c$ as a fitting parameter and then evaluated whether it was smaller or larger than $\delta$, as in similarly done in (Fieremans et al., 2016). In our results, the fitted $t_c$ values ranged from 1 to 3 $\mu m$, which is smaller than $\delta$, justifying the choice $\tilde{t}_c = \delta$.

Assuming the short-range disorder ($p = 0$) and $\tilde{t}_c = \delta$, we obtain (Lee et al., 2020b):

$$D(t) = \begin{cases} D_\infty + 2A_1\, t^{-1/2}, & d = 1 \Rightarrow \theta = \frac{1}{2} \\ D_\infty + A_2\, \dfrac{\ln(t/\delta)}{t}, & d = 2 \Rightarrow \theta = 1 \end{cases} \tag{3}$$

$$K(t) = \begin{cases} K_\infty + \dfrac{4A_1}{D_\infty}\, t^{-1/2}, & d = 1 \Rightarrow \theta = \frac{1}{2} \\ K_\infty + \dfrac{6A_2}{D_\infty}\, \dfrac{\ln(t/\delta)}{t}, & d = 2 \Rightarrow \theta = 1 \end{cases} \tag{4}$$

These expressions provide a theoretical framework for interpreting the time dependence of diffusion and kurtosis. By analysing $D(t)$ and $K(t)$, one can infer key structural properties, such as restriction strength and long-range spatial correlations within the medium. In the context of numerically generated WM substrates, if the synthetic diffusion MRI data follows the predicted time-dependent behaviour of $D(t)$ and $K(t)$ as described by the structural disorder model, this indicates that the substrate realistically captures the microstructural complexity found in actual brain tissue. Agreement with the model thus serves as complement to the validation of the realism of the simulated architecture in this work.

#### 3. Structural correlations within numerical substrates

This complementary analysis builds upon the findings of Figure 10, which examined the time dependence of diffusion and kurtosis under the short-range disorder framework. Here, we compute the two-point correlation function $\Gamma(\mathbf{k})$ for the beaded and undulating substrate $S3$. This function, derived directly from the structure of the medium without the need for MC simulations, represents the power spectrum of spatial restrictions within the substrate. Since our earlier results indicated that $S3$ exhibits short-range disorder, we expect $\Gamma(\mathbf{k})$ to plateau at low spatial frequencies $k_c$, indicating a lack of long-range spatial correlations beyond a characteristic length scale $l_c$. Measuring $\Gamma(\mathbf{k})$ therefore allows us to (i) further confirm that $S3$ belongs to the short-range disorder class and (ii) estimate $l_c$, which was not directly accessible in the previous analysis.

To obtain $\Gamma(\mathbf{k})$, we generated a 3D binary mask $\varrho$ that samples the volume of the numerical substrate of interest at an isotropic resolution of 0.3 $\mu m$. The two-point correlator was calculated from the Fourier transform of the mask using the Wiener-Khinchin theorem $\Gamma(\mathbf{k}) = |\varrho(\mathbf{k})|^2 / V$, where $V$ is the substrate volume. We estimated the two-point correlation function $\Gamma$ in the $x$-$y$ plane by averaging $\Gamma$ along both axes to obtain $\Gamma_{xy}$ (Figure 3A and B), which reflects the structural organisation perpendicular to the axons and along the $z$-axis (Figure 3C and D), representing the primary axonal orientation.

Both two-point correlation functions in the plane perpendicular to the axon and along the axons indeed showed a plateau for frequencies below $k_c \approx 1/10\mu m$, as hypothesised. We therefore confirm the expected short-range disorder, with correlation lengths exceeding $l_c \approx 10\mu m$. This value for $l_c$ falls within the expected range 1-20$\mu m$ reported in (Fieremans et al., 2016).

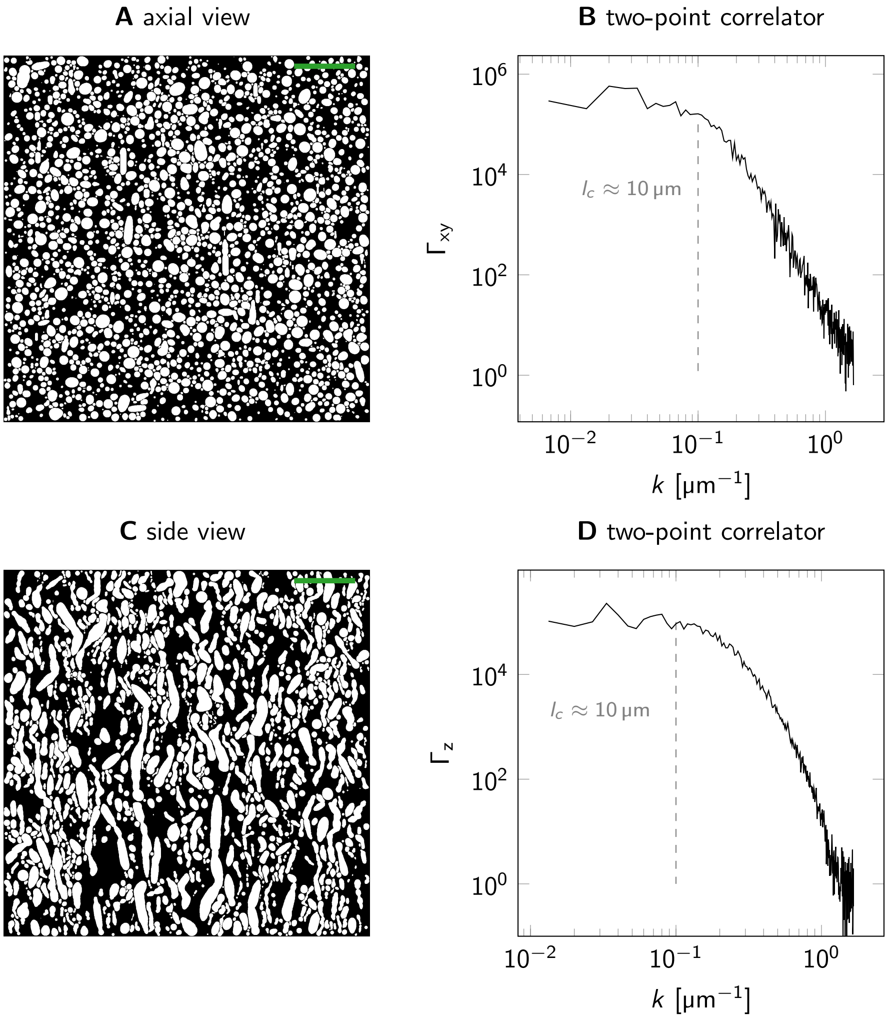


Figure 3: A) Axial view illustrating synthetic axons within a voxel with a volume fraction $f = 50\%$. B) The two-point correlation function $\Gamma_{xy}$ in the transverse plane of the generated voxel. Below a spatial frequency of $k_c = \frac{1}{10\mu m}$, a plateau is observed, which corresponds to short-range disorder for lengths longer than the correlation length $l_c \approx 10\mu m$. C) Lateral (side) view of the same voxel. D) The two-point correlation function $\Gamma_z$ computed along the z-axis, i.e., the primary direction of the axons. As in B, short-range disorder is observed with a similar correlation length of $l_c \approx 10\mu m$. The scale bar in A and C corresponds to $25\mu m$.
